# *Drosophila* Atlastin regulates synaptic vesicle mobilization independent of Bone Morphogenetic Protein signaling

**DOI:** 10.1101/2022.09.09.507308

**Authors:** Francisca Bertin, Jorge Jara-Wilde, Benedikt Auer, Andrés Köhler-Solís, Carolina González-Silva, Ulrich Thomas, Jimena Sierralta

## Abstract

Motor neurons are highly dependent on membrane trafficking, in which the endoplasmic reticulum (ER) and its contact sites with endosomes, confer the ER the role of a long-distance communicator. Atlastin (Atl), a large GTPase located on the ER membrane is required for its function and its tubular structural dynamics. Atl also downregulates, by a yet unknown mechanism, the BMP (Bone Morphogenic Protein) pathway. In humans, Atl mutations are the second more common cause of Hereditary Spastic Paraplegia (HSP), a genetic disease characterized by spasticity of the lower extremities. Here, we explore the molecular basis of Atl-dependent defects on synaptic vesicle (SV) traffic in *Drosophila* under the hypothesis that those defects are the direct consequence of the *atl*-knock-down and not of the Atl-dependent BMP signaling upregulation.

Motor neuronal knockdown of *atl* (Atl-KD) leads to an increase in synaptic and satellite bouton number similar to the increase in BMP signaling activity (TKV-CA). Neuronal Atl-KD also associates to a reduction in the boutons of the abundance of the SV markers CSP (Cysteine string protein) and VGLUT (vesicular glutamate transporter) as well as in TKV-CA larvae, both phenotypes are suppressed by decreasing the function of BMP receptor wishful thinking expressing one copy of the mutant receptor (wit /+). Surprisingly, we determined in Atl-KD larvae an increase in the CSP peripheral density and distribution, dependent on synaptic stimulation, that was not replicated in Tkv-CA larvae, suggesting that there could be differences in the mechanisms that underlie the reduction in CSP abundance. Additionally, we determined that Atl-KD associates to an increase in FM 1-43 unload but not in TKV-CA larvae. Moreover, one copy of *w*it was not able to suppress the FM-143 in Atl-KD larvae (Atl-KD, wit), supporting that BMP signaling does not participate in this phenotype. Together with the stimuli-dependent changes in the SV distribution and dynamics determined in Atl-KD larvae, we measured an increase in Rab11/CSP colocalization, suggesting changes in SV traffic through late recycling endosomes. Together our results suggest a mechanism by which the loss of an ER structuring protein in the motor neuron could, through its role in regulating SV and endosomal trafficking, explain defects in SV accumulation and synaptic dysfunction.

## INTRODUCTION

Every voluntary movement in any organism requires communication between the higher motor centers and the musculoskeletal system, where motor neurons are the effectors of motor activity (Kohsaka et al., 2012). Development and physiology of these polarized and secretory cells depend on the membrane traffic process, in which Rab GTPase proteins exert an essential role (Chan et al., 2011; Raiborg et al., 2015). These proteins are essential for vesicle traffic as well as for vesicle and endosome biogenesis through their role in membrane fusion (Stenmark, 2009). These functions, fundamental for cellular maintenance, are also required for synaptic communication and signaling pathways (Delcroix et al., 2003; McCaffrey et al., 2001; Stenmark and Olkkonen, 2001).

The extended presence of the endoplasmic reticulum (ER) in neurons, from the nuclear envelope to the plasma membrane reaching into the axonal presynaptic terminal, as well as its close communication with multiple organelles such as endosomes, gives this structure a role as long-distance intracellular connector and regulator of signaling (English and Voeltz, 2013; Mattson et al., 2000; Rowland et al., 2014). These ER-mediated functions require the structural flexibility and the regulation of the contact network dynamic, in which ER-organizing proteins such as Atlastin (Atl) are fundamental (Park et al., 2010). Atl is a large GTPase protein member of the dynamin superfamily; the 3 mammalian genes are widely expressed, of which Atl-1 is mainly expressed in the nervous system where it localizes to the ER, Golgi apparatus and endosomes (Park et al., 2010; Rismanchi et al., 2008). Atl displays a homotypic membrane fusion function, which confers the characteristic orthogonal conformation of the axonal tubular ER (Hu et al., 2015; Rismanchi et al., 2008), and it regulates morphogenesis and dynamic structural changes of the Golgi apparatus and ER. In endosomes, however, its function is less known, but it has been suggested to exert a regulatory role in membrane traffic (Liu et. al., 2019). In *Drosophila*, only one atlastin (*atl*) gene has been described, and a role in trans synaptic signaling modulation, specifically a downregulation of BMP (Bone Morphogenic Protein) signaling has been reported (Fassier et al., 2010; Summerville et al., 2016). BMP signaling at the neuromuscular junction (NMJ) is a homeostatic signal that involves the retrograde signaling by the muscle-derived ligand Gbb (Glass Bottom Boat) onto serine/threonine kinase receptors Wit (Wishful Thinking), Tkv (Thickveins) and Sax (Saxophone), in the motor neuron. (Deshpande and Rodal, 2016; Frank et al., 2020; McCabe et al., 2003). The activation of the receptors promotes the phosphorylation of the signaling complex (Wit/Tkv or Wit/Sax), which is endocytosed and trafficked retrogradely in signaling endosomes to the soma, where these activated receptors phosphorylate the cytosolic protein MAD (Mothers Against Decapentaplegic, homologous to mammalian SMAD), pMAD translocate to the nucleus, promoting transcription of genes such as *trio* (Ball et al., 2010; McCabe et al., 2004; Smith et al., 2012). Also, in the terminal, a local phosphorylated MAD present in the synaptic bouton can be found, which is involved in synaptic development independent of the nuclear signal (Smith et al., 2012; Sulkowski et al., 2016). The BMP signaling involves modifications in the actin cytoskeleton and microtubules, its hyperactivation promotes the formation of satellite boutons, corresponding to small ectopic boutons that emerge from pre-existing synaptic boutons or from the main terminal branch of the NMJ (O’Connor-Giles et al., 2008; Wang et al., 2007). Atl inhibits BMP pathway signaling in primary culture of zebrafish spinal motor neurons, colocalizing with the Rab GTPase proteins: Rab4 (regulator of the rapidly recycling endosome) Rab5 (regulator of the early endosome), and Rab7 (a regulator of the late endosome) as well as with BMP receptors (Fassier et al., 2010). In *Drosophila*, an interactome study determined that Atl interacts with Rab4, which is also involved in the rapid recycling of endocytosed components, mediating the redistribution of cargo from early endosomes to the plasma membrane, towards the Rab11-positive late recycling endosomes or to Rab7-positive late degradation compartments (McCaffrey et al., 2001; O’Sullivan et al., 2013). However, a molecular mechanism that integrates the antagonistic role of Atl on BMP signaling, and its relationship to the endocytic pathway is unknown (Fassier et al., 2010; O’Connor-Giles and Ganetzky 2008; Wang et al., 2007).

Alterations in ER morphology and function are associated with defects in fundamental physiological processes of neurons, such as calcium homeostasis and membrane trafficking. These alterations have been associated with the development of several neurodegenerative diseases, such as Huntington’s disease, Amyotrophic Lateral Sclerosis, and Hereditary Spastic Paraplegia (HSP) (Kegel et al., 2000; Namekawa et al., 2006; Squitieri et al., 2010; Vollrath et al., 2014). In HSPs, Atl autosomal dominant mutations are associated in 10% of the cases (Blackstone, 2018).

In *Drosophila, atl* null mutations lead to progressive motor defects and the morphology of larval NMJs shows an increase in satellite bouton numbers and ER structural abnormalities. These phenotypes were accompanied by an increase in motor neuronal BMP signaling as reflected by elevated nuclear pMAD (De Gregorio et al., 2017; Lee et al., 2009; Summerville et al., 2016). Also, Atl-KD larvae exhibit an accumulation of synaptic vesicle (SV) and lysosomes markers in distal axons, yet a reduced SV number in the nonadjacent area surrounding the active zone, as well as defects in the recovery of synaptic function after a tetanic stimulus (De Gregorio et al., 2017). It has been described those mutations in endosomal proteins repressors of BMP signaling (implying an increase in this signaling), generate a reduction in SV number, modifying the number and size of those vesicles located within the active zone, a phenotype not observed in Atl knockdown larvae. On the other hand, in larvae with increased BMP signaling in motor neurons, the axonal accumulation of SVs as detected in Atl knockdown larvae was not observed (Shi et al., 2013; X. Wang et al., 2007; G. Zhao et al., 2015). These discrepancies in phenotypes suggest differences in the mechanisms underlying BMP signaling and Atl functions, respectively. We hypothesized that SV defects are a direct result of Atl knockdown, associated with alterations in the SV intracellular trafficking, and not a downstream consequence of increased BMP signaling. In support of this hypothesis, our results suggest a new mechanism by which loss of function of an ER structuring protein in the motor neuron could, through its role in regulating endosomal trafficking, explain defects in SV accumulation, even allowing us to establish that defects in membrane trafficking are part of the pathological mechanism underlying the distal axonopathy present in Atl-determined HSP.

## RESULTS

### *atl* knockdown in motor neurons increases BMP signaling

In *Drosophila, atl* null mutant larvae display an increase in neuronal nuclear pMAD accumulation, implying an increase in BMP signaling, as well as an increase in the number of synaptic boutons, a phenotype associated with an enhancement in BMP signaling (De Gregorio et al., 2017; Summerville et al., 2016). To confirm that the phenotype observed in motor neurons is due to an increased neuronal BMP signaling, we determined synaptic pMAD accumulation in larvae with neuronal *atl* knockdown by Gal4-driven RNAi (Atl-KD) (Figure 1A-E). Also, in this work, we focused our analyses on abdominal segment A6, where the motor neuronal axons are longest and thus particularly prone to distal axonopathy (DeGregorio et al. 2017). As positive control, we evaluated synaptic pMAD accumulation in larvae with an overexpression of the constitutively active Tkv receptor (Tkv-CA). Both *atl*-KD and Tkv-CA expression were carried out using two different motor neuronal promoters, C380-GAL4 and OK6-GAL4. Tkv-CA expression with either Gal4 driver induced a significant increase in synaptic pMAD (33% and 56%, respectively, Figure 1B-C). A similar increase in synaptic pMAD was observed in Atl-KD larvae (30% for C380-GAL4 promoter, and 47%, for OK6-GAL4, Figure 1B-C; Suppl Figure 1). These results demonstrate that the observed increase in neuronal BMP signaling in *atl* mutants can be replicated with motor neuron specific Atl-KD. To confirm the specificity of this phenotype, we determined the synaptic pMAD accumulation in Atl-KD larvae with a decreased allelic dose of the Wit receptor using a *wit* null mutant allele (Atl-KD/*wit*). While pMAD levels in heterozygous *wit* /+ larvae showed a 31% reduction, heterozygosity for this mutation also precluded the increase of synaptic pMAD levels in Atl-KD larvae (Figure 1D-E, Supp. Figure 2). Thus, we show here that neuronal Atl-KD replicates *atl* mutant BMP activation phenotype, which can be rescued by decreasing the dose of BMP-receptors.

**Figure 1:**
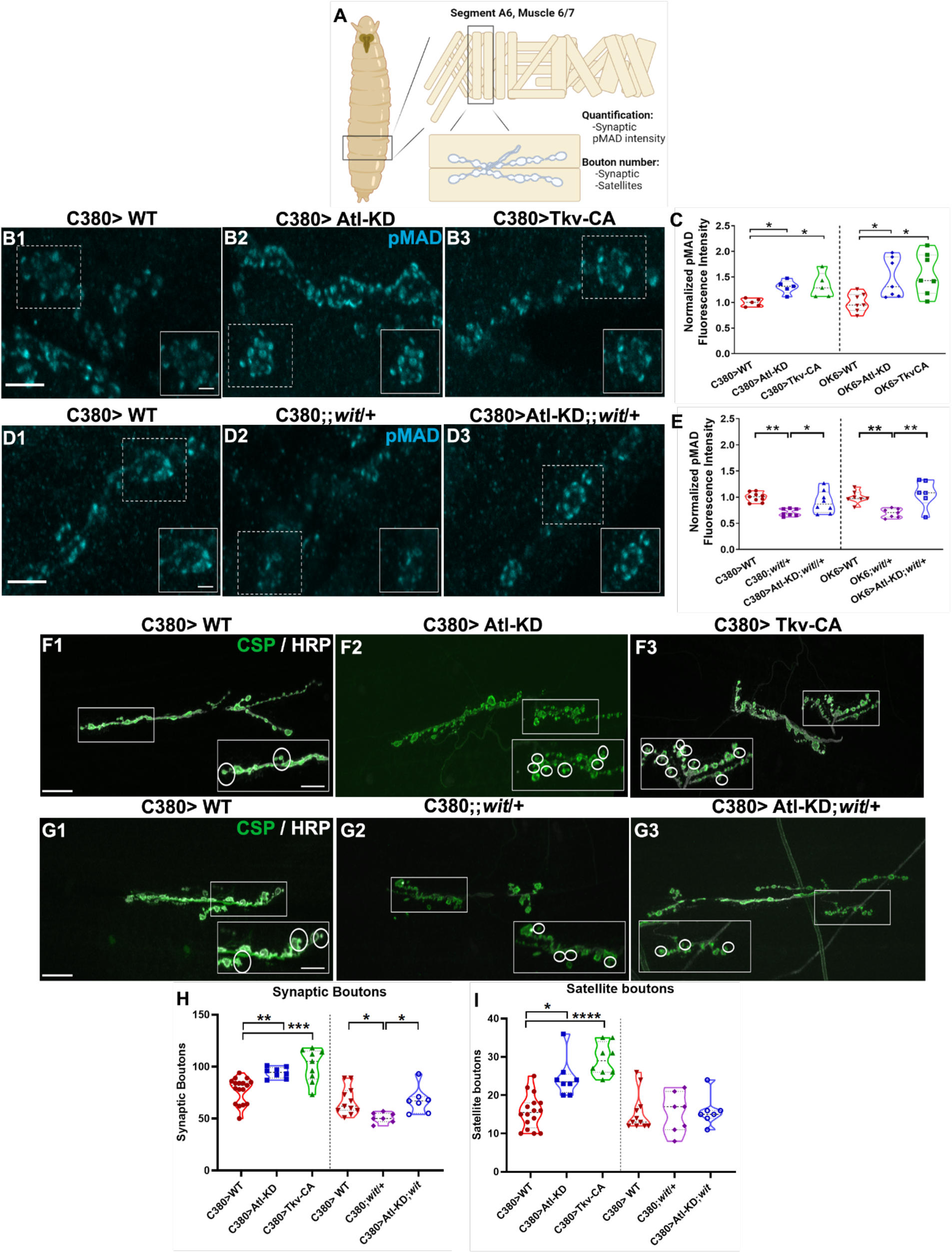
*atl* knock-down in motoneurons increases synaptic pMAD and morphometric parameters. A) *Drosophila* larvae NMJ were utilized to observe alterations in Atl-KD, Constitutively Tkv-CA and wit/+. C380-GAL4 was used to overexpress UAS constructs on motoneurons. Cartoon created with BioRender.com. B, D) Confocal microscopy images (maximum intensity Z-projection) of synaptic pMAD of control (C380), Atl-KD, TkvCA, *wit* and Atl-KD;*wit* larvae. pMAD antibody stain labeled in cyan color. Scale bar of large image: 5μm, of cropped image: 2μm. C) Synaptic pMAD intensity of C380, OK6 Atl-KD and TkvCA larvae: pMAD intensity normalized to control levels. E) Synaptic pMAD of C380, Ok6; *wit* and Atl-KD;*wit* larvae: pMAD intensity normalized to control levels. F-G) Confocal microscopy images (maximum intensity Z-projection) of presynaptic terminal of C380, Atl-KD, TkvCA, *wit* and Atl-KD;*wit* larvae. CSP and HRP antibody stains labeled in green and gray color, respectively. In cropped images, satellite boutons are shown within white circles. Image intensity and contrast have been increased for visualization purposes. Scale bar of large image: 20μm, of cropped image: 10μm. E-F) Synaptic and satellite bouton number of C380, Atl-KD, TkvCA larvae motor neuron. H-I) Synaptic and satellite bouton number of C380, *wit* and Atl-KD; *wit* larvae. Each scatter dot represents one measurement. Kruskal-Wallis, p-value *<0,05; **<0.01; ***<0.001. n=5-9.

### *atl* knockdown and BMP activation in motor neuron increases synaptic and satellite bouton number

In *Drosophila*, altered synaptic function at larval NMJs is often found associated with variations in neuronal morphology such as the occurrence of so-called satellite boutons. Satellite boutons refer to small supernumerary boutons that emerge from a parental bouton, indicating an immature bouton state (O’Connor-Giles and Ganetzky, 2008). Loss of *atl* function generates an increased number of satellite boutons, along with deficiencies in synaptic function (De Gregorio et al., 2017; Summerville et al., 2016). An increase in satellite bouton number has also been associated with BMP activation, related to changes in the organization of the actin and microtubule cytoskeleton (Ball et al., 2010; Pawson et al., 2008). However, in various endocytic mutants, where BMP signaling modifications have not been reported, such an increase was also described, suggesting that these boutons could emerge as a compensatory response to endocytic failure (Dickman et al., 2006). Given these differences, we aimed to determine the contribution of BMP signaling to the atl-KD NMJ structural phenotype. For this, we quantified and compared the number of mature and satellite boutons at both Atl-KD and Tkv-CA NMJs. C380-driven Atl-KD significantly increased the number of synaptic boutons (47%, Figure 1F-H), similarly as the expression of Tkv-CA (31%, Figure 1F-H). The number of satellite boutons present in Atl-KD larvae was increased to 66% of the total bouton number, similar to Tkv-CA larvae (61%, Figure 1F-F’’,I). Very similar findings were made, when OK6-Gal4 was used instead of C380-Gal4 (Suppl. Figure 1).

To evaluate BMP contribution to these morphometric parameters, we counted mature and satellite boutons at NMJs of *wit* /+ larvae and for atl-KD NMJs in a *wit*/+ background (Atl-KD/ *wit*). We found that the *wit* /+ mutation is sufficient to significantly reduce the number of synaptic boutons by 24%, while also efficiently precluding the increase of synaptic boutons in Atl-KD larvae (Figure 1G-H). Regarding the number of satellites, the *wit* /+ mutants and controls were indistinguishable. However, a decreased dose of *wit* was efficient in preventing the satellites increase in *atl*-KD larvae. (Figure 1G-G’’,I). This morphometric characterization establishes that both Atl knockdown and BMP modifications alter synaptic and satellite bouton number in the motor neuron. Furthermore, a genetic decrease in BMP signaling in Atl-KD larvae is sufficient to prevent the increased synaptic and satellite boutons number observed in these larvae. This supports that Atl knockdown induces the BMP activation, which in turn leads to an increased abundance of both mature and satellite boutons.

### *atl* knockdown increases CSP peripheral density after stimulation

Using electron microscopy, De Gregorio et al. (2017) determined that Atl knockdown in motor neurons reduced the number of synaptic vesicles (SV), in a localized manner. At the same time, a striking axonal accumulation of the SV marker CSP, which colocalized with the lysosomal marker Lamp2, was observed. This axonal phenotype has not been described for *Drosophila* larvae that overexpress the Tkv-CA receptor (Wang et al., 2007). To further address this phenotype and the impact of BMP signaling onto it, as well as on the SV recycling at the axon terminal, we used antibodies against the SV proteins CSP and dVGLUT, in both Atl-KD and Tkv-CA larvae. Atl knockdown and Tkv-CA expression in motor neurons both significantly reduced CSP abundance in the synaptic boutons (Supp.Fig S3A). Consistently, the abundance of the dVGLUT marker also was decreased significantly in both genotypes (Figure not shown). To determine the contribution of BMP to this phenotype, CSP was also quantified in Atl-KD in *wit/+* background. Heterozygous *wit* mutation alone did not show phenotype, however as a genetic background it precluded the Atl-KD-induced reduction of CSP (Figure2A’, Supp.Figure 2). Thus, both Atl knockdown and BMP activation in motor neurons reduced the accumulation of synaptic vesicle markers on the bouton. On the other hand, the reduction in BMP signaling caused by the *wit* /+ mutation, in Atl-KD larvae, prevented the CSP decrease suggesting that the BMP signaling contributes to the SV phenotype observed in Atl-KD larvae. The SV number reduction located not adjacent to the active zone determined by Gregorio et al. (2017) together with the deficiencies in synaptic function recovery after a prolonged tetanus stimulus suggests that Atl knockdown modifies intracellular traffic. Moreover, modifications in the size and abundance of SV have also been described in endosomal protein mutants that modulate the BMP pathway, suggesting that this signaling could be involved in this phenotype (Dickman et al., 2006; Shi et al., 2013; G. Zhao et al., 2015). To determine if the defects observed in CSP abundance are also observed associated with modifications in synaptic activity, we analyzed the density distribution of CSP markers in both Atl-KD and Tkv-CA larvae, before (**US**, unstimulated) and after synaptic stimulation (90 mM KCl, **S**, stimulated) using super resolution microscopy STED (Stimulated Emission Depletion) (Figure 2B). CSP distribution analysis was performed in the periphery (first 200 nm from the border, associated to the readily releasable pool of SV) and in the center of the bouton (associated to the reserve pool).

**Figure 2:**
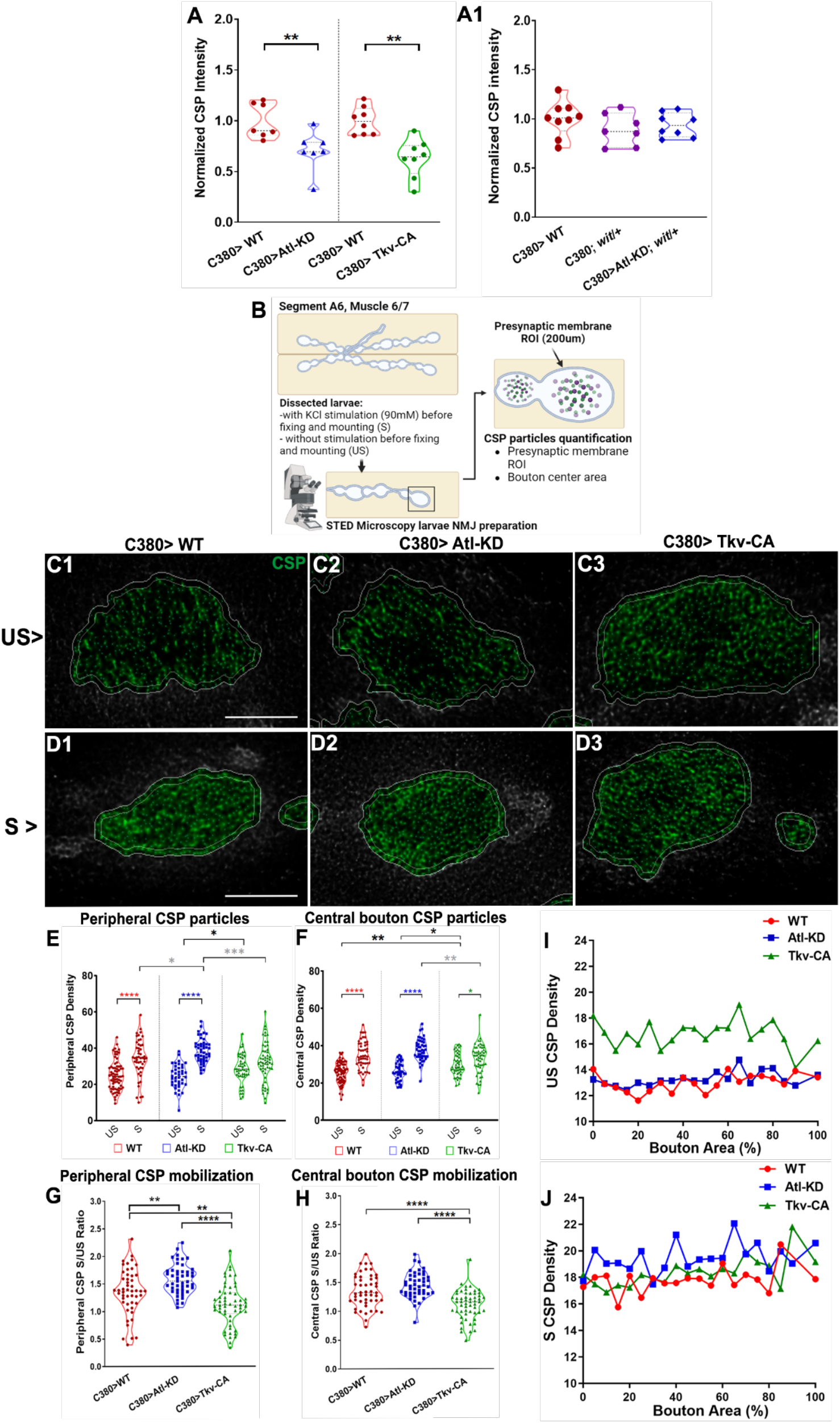
Synaptic stimulation increases peripheral CSP density in *atl* knock-down larvae. A) Normalized CSP intensity of WT, Atl-KD and TkvCA larvae. Each scatter dot represents one measurement. C380>WT vs Atl-KD, Mann-Whitney p-value ** <0.01; C380>WT vs Tkv-CA, Mann-Whitney p-value ** <0.01. For WT, wit/+ and Atl-KD; wit/+ analysis an ANOVA Kruskal-Wallis analysis was performed. n=7-9. B) Acquiring STED images from WT, Atl-KD and TkvCA *Drosophila* NMJ, with (S) or without (US) KCl Stimulation; larvae were fixed and processed for immunostaining after the stimulation or non-stimulation protocol. Each particle set was quantified in the peripheral delimited area (ROI generated from the HRP), as well as in the center of the bouton. Cartoon created with BioRender.com. C-D) Representative images of peripheral and central CSP density in synaptic bouton of control (C380), Atl-KD and TkvCA larvae. Scale bar: 2μm. E-H) Peripheral and central CSP density of C380, Atl-KD and TkvCA larvae: CSP density in unstimulated (US) and stimulated (S) larvae motor neurons. F-G) CSP mobilization of C380, Atl-KD and Tkv-CA larvae: CSP density S/US ratio of larvae motor neurons. I-J) CSP density distribution on the whole synaptic bouton, under non-stimulated (US) and stimulated conditions, for C380, Atl-KD and Tkv-CA conditions. Each scatter dot represents one measurement. One-way ANOVA, p-value *<0,05; ** <0.01;***<0.001; ****<0.0001. n=5 larvae.

In this analysis, we included only synaptic boutons between 2.5-12 μm^2^ (type Ib, size distribution observed). Comparing CSP density in the periphery of the bouton, before and after the stimulus within each group, control larvae displayed a significant increase between unstimulated and stimulated conditions, showing that the CSP labeling was able to detect the mobilization of SV after synaptic stimulation by KCl (Figure 2C, D). In the Atl-KD group, KCl stimulation triggered, as in control larva, a significant increase in the peripheral CSP labeling (Figure 2E). This increase was, however, significantly larger than the observed in the control. Notably, and contrary to the phenotypes observed previously, a peripheral density variation was not significant in Tkv-CA larvae (Figure 2C-E), suggesting that the BMP signal affected the SV mobilization downstream of Atl. The analysis in the center of the boutons, showed again a significant increase in the CSP density in control larvae, which suggests that the reserve compartment increased after the synaptic activity induced by the potassium stimulus (Figure 2F). In Atl-KD larvae, CSP signal density increased similarly to the control group, showing no difference with the control group in its basal levels or the stimulated condition (Figure 2F). However, Tkv-CA larvae already showed an increase in the central density of CSP in basal conditions (prior to stimulation) compared to control and Atl-KD and only a moderate though significant increase after stimulus (Figure 2F). We therefore compared the ratio of CSP density change between US and S conditions. for both the periphery and the center of boutons. The ratio of density changes in the periphery of the synaptic bouton in Atl-KD group displayed increased peripheral redistribution, compared to the genetic control (Figure 2G). On the contrary, Tkv-CA larvae even showed a slightly reduced peripheral distribution by 17%, compared to control (Figure 2G). Tkv-CA larvae also showed a reduction in the center of boutons by 18% with respect to control (Figure 2H). Given that this analysis focused on studying the changes in the periphery of the bouton, leaving out the innermost regions, and thus obscuring variations in the general organization of SV within it, we conducted a complementary analysis. In this analysis, all the synaptic boutons present in the images were included. The cumulative CSP density data was organized into fixed bouton percentage area segments (5%), allowing a comparative area percentage analysis for various bouton sizes. Qualitatively, the accumulated CSP density for each percentage of bouton area was plotted, from the center of the bouton (0%) to the periphery (100%, total area), both before and after the stimulus (Figure 2I-J). Using this analysis, the CSP densities present in each percent quartile of the bouton were quantified, comparing these values between each genotype and experimental condition (Figure 2I-J). Tkv-CA larvae presented an increased accumulated density of CSP in 29%, while Atl-KD larvae showed, after the stimulus, an increased CSP density in 9%, in reference to control. Again, it was established that the Tkv-CA larvae, but not the Atl-KD, displayed an increased CSP density within each quartile of the bouton, in basal conditions. On the contrary, and consistent with previous results, Atl knockdown generates an increase in CSP density mainly restricted to the peripheral regions of the bouton, being significant in the last 2 quartiles (75 and 100) and dependent on neuronal stimulation. A significant increase in CSP density was also determined in the center of the bouton, which suggests alterations in CSP redistribution after stimulation.

### *atl* knockdown in motor neurons modifies synaptic vesicle dynamics and distribution through recycle components in *Drosophila* NMJ

To confirm the presented findings, we evaluated SV dynamics (endocytosis and exocytosis) using the lipophilic dye FM1-43, in which the discharge (unloading) of the previously endocytosed dye (loading), reports the SV mobilization (Gaffield and Betz 2006, Figure 3). For this analysis, motor neurons were stimulated chemically (90 mM KCl). Dissected larvae were incubated for 3 minutes with FM1-43 in HL3.1 solution and 90 mM KCl, to promote the exocytosis of SV and subsequent membrane recycling, where each mobilized vesicle will capture within their membranes the FM1-43 available in the medium. Subsequently, medium washing removed the excess of this product from the outer membrane. Then, larvae were photographed in the microscope in medium without calcium and with 0.5mM EGTA, this corresponded to the loading image of FM 1-43. Then, a second stimulation was carried out, again using the previously described KCl solution, but in the absence of dye, promoting exocytosis of the labeled vesicles. For the acquisition of discharge images, the larvae were washed and kept in HL3.1 with 0.5mM EGTA (Figure 3A). In this analysis, the discharge index corresponds to the average fluorescence of each synaptic bouton obtained from the image, normalized by the average load of the bouton. The discharge percentage corresponds to the value 1 minus the previously mentioned discharge index. Atl knock-down in motor neurons does not significantly modify the load of FM 1-43, regardless of the expression promoter used, however, these larvae displayed increased FM 1-43 unloading, compared to control (Figure 3D-E, Supp.Figure3). On the other hand, the analysis of SV dynamics in larvae with *wit*/+ mutation and Atl-KD, *wit*/+ line showed that, like control groups, *wit*/+ mutation did not modify the load of the dye or its discharge (Figure 3H-I). However, Atl-KD; *wit* larvae, like the Atl-KD group alone, has an increased discharge in respect to the wit larvae, suggesting that the BMP signal is not involved in the SV phenotype (Figure 3I).

**Figure 3:**
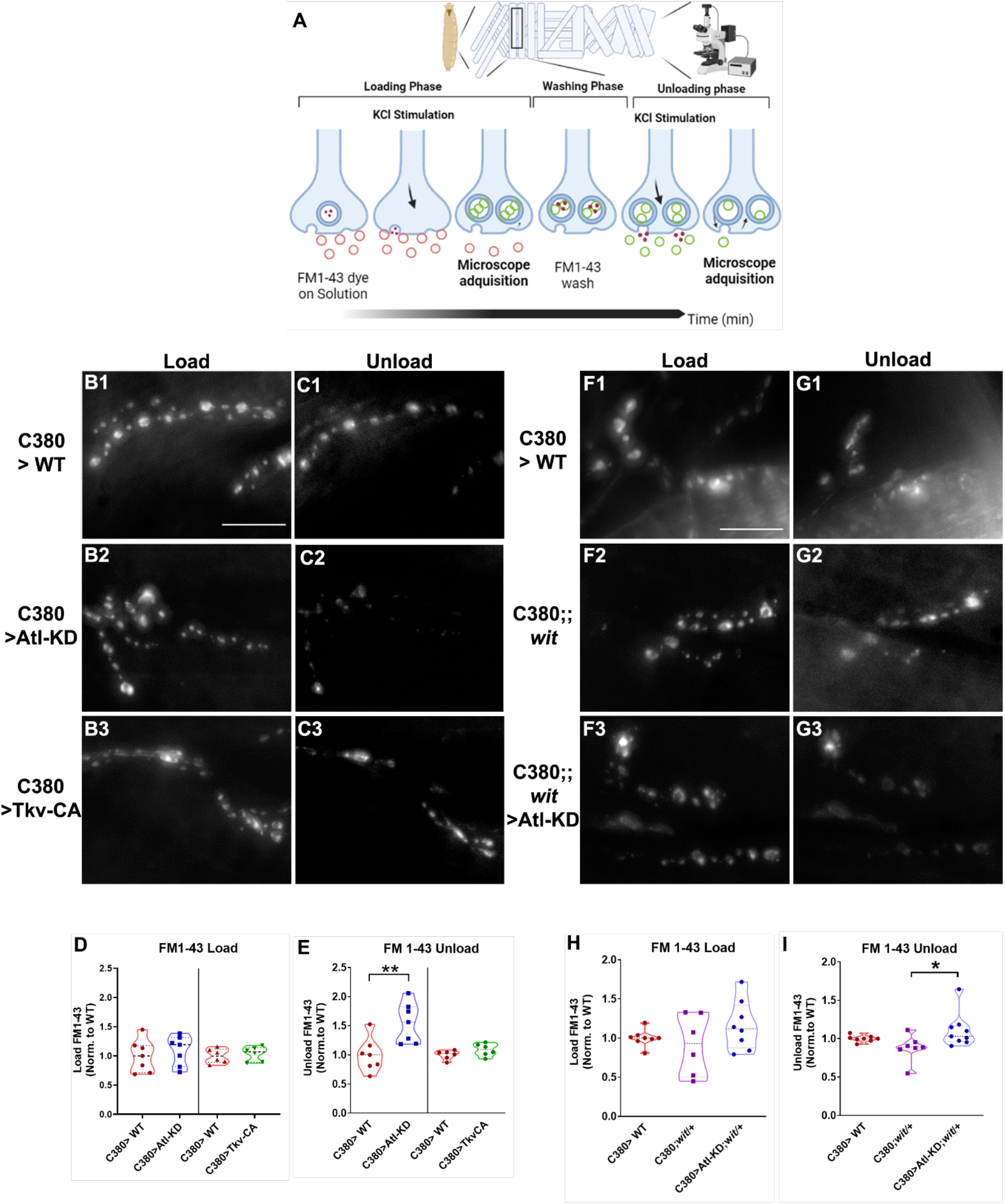
*atl* knock-down in motor neuron increases FM 1-43 unloading. A) FM1-43 technique on *Drosophila* larvae NMJ. Cartoon created with BioRender.com B-G) Spinning disc images (maximum intensity Z projection) of FM1-43 load and unload in the presynaptic terminals of control (C380) (B-C, F-G), Atl-KD (B’-C’), constitutive active TkvCA (B’’-C’’), *wit* mutant (*wit*) (F’-G’) and Atl-KD;*wit* (F’’-G’’) larvae. Image intensity and contrast have been increased for visualization purposes. Scale bar: 20μm. D-I) FM 1-43 loading in C380, Atl-KD and TkvCA larvae: FM 1-43 load normalized to control levels. E) FM 1-43 unloading in C380, Atl-KD and TkvCA larvae: FM 1-43 unload normalized to control levels. Each scatter dot represents one measurement. H) FM 1-43 load in C380; *wit* and Atl-KD; *wit*: values normalized to control. I) FM 1-43 unload in C380; *wit* and Atl-KD; *wit*: values normalized to control. Each scatter dot represents one measurement Kruskal-Wallis, p-value *<0,05, **<0.01, n= 6-9 larvae.

The differences in the distribution and density change rate of CSP, indicative of mobilization, between both Atl-KD and Tkv-CA larvae, determined before and after the stimulus, show that the mechanisms underlying the reduction in the SV markers abundance in these groups are distinct. Since our results are compatible with a defect in the process of vesicle mobilization or turnover in Atl-KD larvae, we analyzed the colocalization of rab-proteins involved in the process of endocytosis and recycling with synaptic vesicles labeled with CSP, to determine difference between control and Atl-KD group (Figure 4A). We did not detect differences in the Manders 1(M1) and Manders 2(M2) coefficients between Atl-KD and control groups for Rab-7, which labels the late endosome, or Rab-4, which labels early recycle compartments (Figure 4B, C, E, F). However, we did detect a significant difference in M2 for Rab11, a marker for late recycling compartments (Figure 4D, G). M2 represents the colocalization of the Rab11 signal on CSP’s, respecting all RAB11 signals, while M1 describes the colocalization of CSP in RAB 11 considering all CSP signals. As the CSP signal is much more abundant and widely distributed than Rab 11, the M1 is not very informative. However, M2 values suggest that Rab11 is associated with SVs in the Atl-KD group more than in the control group, suggesting that Atl regulates the traffic of SV in the late recycling compartment.

**Figure 4:**
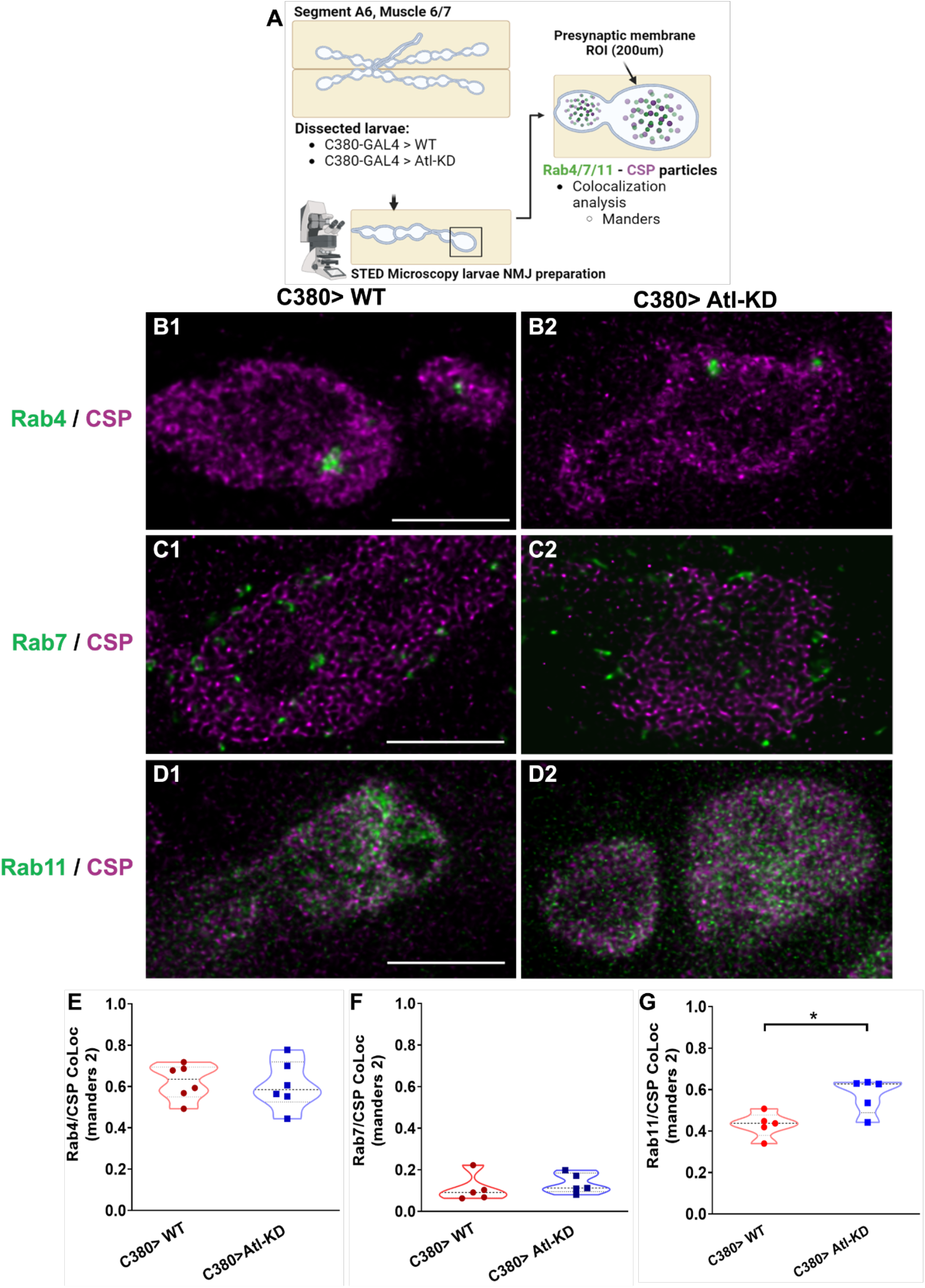
*atl* knock-down in motor neurons increases the colocalization of Rab11 with CSP. A) Acquiring STED images from WT and Atl-KD *Drosophila* NMJ, using Rab4, Rab7 and Rab11 GFP fused UAS constructs on motoneurons. Cartoon created with BioRender.com. B-D) STED microscopy images (maximum intensity Z projection) of the presynaptic terminal of control (C380) and Atl-KD larvae expressing a UAS-Rab4-GFP. Scale bar: 2μm (B), endogenous Rab7-GFP (C) and UAS-Rab11-GFP (D). CSP antibody stain is represented in Magenta. Image intensity and contrast have been increased for visualization purposes. Scale bar: 2μm. E-G) Rab4/CSP colocalization: Manders 2 coefficient of Rab4 signal colocalized with CSP marker, divided total Rab4 signal. F) Rab7/CSP colocalization: Manders 2 coefficients of Rab7 signal colocalized with CSP marker, divided total Rab7 signal. G) Rab11/CSP colocalization: Manders 2 coefficient of Rab11 signal colocalized with CSP marker, divided total Rab11 signal. Each scatter dot represents one measurement. Mann-Whitney, p-value *<0,05, n=5-6 larvae.

## DISCUSSION

In the *Drosophila* NMJ and in other model systems, loss of Atl function has been associated with an increase in BMP signal (Fassier et al., 2010; Summerville et al., 2016). Thus, the question that we sought to answer was whether the synaptic *atl* phenotype exclusively results from exacerbation of the BMP pathway, or alternatively, is only partially attributable to defects in this pathway. In favor of the latter alternative, it has been reported that Atl colocalizes and interacts with various Rab proteins, and a reduction in the number of SV not adjacent to the active zone, suggesting that intracellular transport of recycled vesicles is affected in *atl* mutant animals (De Gregorio et al., 2017; O’Sullivan et al., 2013). In this work we were able to establish that motor neuronal knockdown of Atl is sufficient to significantly increase synaptic pMAD accumulation, as well as the number of synaptic boutons and satellite boutons at NMJs, and both phenotypes are associated with increased BMP signal (Lee et al., 2009; Summerville et al., 2016). However, increases in the number of satellite boutons have also been reported in endosomal mutants not involved in the negative regulation of BMP, and in hypomorphic mutants of calcium-activated potassium channels, therefore the mechanisms underlying the changes in morphometric parameters in *atl* loss of function may not solely be the result of BMP-signal increase (Dickman et al., 2006; Lee and Wu, 2010). Our results show that reduced expression of the BMP wit receptor in the Atl-KD larvae, compensates for the changes in these morphometric parameters (Figure 1), supporting a role for Atl-induced BMP-signal increase in the origin of the morphological phenotype. Another phenotype described by De Gregorio et al. (2017) was the localized reduction in the number of SV not adjacent to the active zone. It is interesting that mutations that generate an overactivation of the BMP pathway also cause a reduction in the number of SV, but mainly adjacent to the active zone (Shi et al., 2013; Zhao et al., 2015). We determined that Atl knock-down reduces the abundance of SV CSP and VGLUT markers (Figure 2A), suggesting that the drop in the number of SVs recorded by De Gregorio et al., (2017) does not correspond to a localized finding, but to a generalized phenomenon within the synaptic bouton, where Atl would be promoting the accumulation of SV in the synaptic bouton. On the other hand, the reduced abundance of CSP in larvae that overexpress the Tkv-CA receptor, like that of Atl-KD larvae, would suggest that the increase in BMP signaling could underlie this phenotype. However, Wang et al. (2007) showed that the overexpression of Tkv-CA, in motor neurons, was not associated with accumulations of SV and lysosomal markers in the axon as described by De Gregorio et al. (2017). This suggests that *atl* gene silencing and increased BMP signaling are sufficient to reduce SV marker levels; however, the mechanisms underlying these accumulation defects could be different. It is interesting that, although the abundance of SV markers was reduced in both Atl-KD and Tkv-CA larvae, the distribution of this marker and its dependence on synaptic activity was different between the two genotypes. Whereas the overactivation of BMP signaling, in the absence of changes in Atl expression, increases basal CSP levels in the center of the bouton, Atl knock-down increases peripheral CSP density, post-stimulation (Figure 2). This result establishes that in the two genotypes that increase BMP signaling there are changes in the organization of the SV with a decrease in CSP abundance in the bouton, however it also suggests that the phenotype in the two genotypes have different underlying mechanisms, since the phenotypes have diverse characteristics and in some cases opposite.

Regarding mobilization, it has been described that strong and sustained synaptic stimuli increase the SV fraction that make up the exo-endocytic population (detailed below), associated with spatial remodeling and increased migration of these vesicles to the active zone (Rey et al., 2020). In Atl-KD larvae, a greater mobilization of CSP was evident in the periphery of the bouton (Figure 2C’-E) This change in density dependent on synaptic activity indicates an increased recruitment of SV towards peripheral regions of the bouton, which could be a compensatory response to maintain synaptic function. In the case of the changes in basal density found in Tkv-CA larvae, as sustained activation of BMP, could be associated with changes in the actin cytoskeleton, involved in the transport of SV, which is related to modifications in the basal distribution and mobilization of VS (Gramlich and Klyachko, 2017).

At a functional level, the SV population has been divided into various populations or pools: A) the readily releasable pool, which largely represents the SV population adjacent to the active zone; B) the recycling pool, which is mobilized to the active zone after the release of the previous pool and C) The reserve pool, corresponding to the SV population that is only released during intense stimulation. It has been determined that the various functional populations of SV differ in their availability of mobilization dependent on a stimulus (Alabi and Tsien, 2012), where light or moderate synaptic stimuli imply the recruitment of pools ready to be released and recycling, forming the exo-endocytic pool. While the characterization of these pools is functional, to date it has not been possible to structurally differentiate these groups, since the mobilization of SV generates a spatial redistribution and exchange between populations (Alabi and Tsien, 2012; Denker et al., 2009). This implies that the changes in density distribution and SV mobilization determined in Atl-KD larvae and those with Tkv-CA receptor overexpression cannot be attributed to any specific pool. The significant increase in FM 1-43 discharge determined in Atl-KD larvae, but not in TkvCA larvae (Figure 3), indicates that the defect in SV mobilization is independent of an increase in BMP signaling. Modifications in SV traffic, where the main pool mobilized is the exo-endocytic, without adequate recruitment compensation or exchange between the various pools, could underlie the higher FM 1-43 discharge (Ball et al., 2010; Uytterhoeven et al., 2011). This phenotype could also be related to the fact that changes in the density and mobilization of peripheral CSPs, where the active zones are located, are evident after synaptic stimulation.

Regarding BMP signaling, the levels of loading or unloading of FM1-43, comparable to the control, in Tkv-CA larvae and *wit* / + mutants, suggest that this phenotype is not sensitive to modifications in BMP signaling. The fall in the abundance of SV markers, the increase in their density and mobilization to peripheral zones of the bouton, after stimulation and the greater mobilization of the exo-endocytic population evidenced in the Atl-KD larvae, could be related to the reported inability to recover synaptic function after a sustained tetanic stimulus, where the modifications in the intracellular traffic of SVs would be sufficient to respond in early phases of this stimulus, but they would not be able to maintain a sustained compensatory response over time (De Gregorio et al., 2017). Regarding Atl and SV, it is interesting that, in rat brain Atl was identified as one of the GTPases included in the SV proteome (Burré et al., 2006). This, together with the results obtained here, suggests that Atl knockdown modifies the mobilization of VS dependent on synaptic activity and that this protein would be playing a regulatory role in the intracellular traffic of SV. An adequate balance between endocytosis, recycling and degradation processes is essential to maintain synaptic homeostasis and neurotransmission. In the SV cycle, Rab proteins involved in these processes include Rab5, Rab4, Rab11 and Rab7 (Dey et al., 2017; Inoshita et al., 2017; Pavlos and Jahn, 2011). The increase in the formation of multivesicular body-like structures in the synaptic bouton, together with the accumulation of SV and lysosomal markers in the distal axons reported by De Gregorio et al. (2017), suggests that Atl knock-down is related to alterations in membrane trafficking, implying possible changes in the organization and life cycle of the SV (Piccini et al., 2017). Here we determined that Atl knock-down modifies the accumulation of SV in the endosomal compartments positive for Rab11, but not for Rab4 (early recycling endosome) or Rab7 (late endosome) (Figure 4). This implies that in motor neurons, the loss of Atl function selectively affects the traffic of SV through this late recycling endosomes. Also, both Atl and Rab11 have been identified as SV components (Burré et al., 2006), thus Atl could modify the destination or transit of the vesicles to or from this compartment. As a potential mediator of this process, we could name prothrudin, a resident protein of the ER that has been reported to associate to Atl and that is implicated in the symptoms of HSP, which as well presents interaction domains with Rab11 (Chang et al., 2013; Hashimoto et al., 2014). Rab11 is a key presynaptic regulator during strong synaptic stimuli (Kokotos et al., 2018). Here we described that the synaptic phenotypes of increased density and peripheral mobilization of CSP, together with the enhanced discharge of the labeled exo-endocytic pool, are evident after synaptic stimulation suggesting that this change is accentuated after activity. The phenotype observed with Rab11, under baseline conditions, could be related to the increase in the number of multivesicular body-like structures previously observed in Atl-KD larvae, a compartment that also interacts with Rab11 and where this protein performs biogenesis, transport and secretion functions (Blanc and Vidal, 2018; De Gregorio et al., 2017; Savina et al., 2005). Modifications in the area or function of Rab11 have been implicated in several neurological pathologies, including Huntington’s disease and amyotrophic lateral sclerosis, which are not only associated with alterations in synaptic function, but also with variations in BMP signaling. (Akbergenova and Littleton, 2017; Mitra et al., 2019). Regarding Atl knock-down, endosomal recycling and BMP signaling, it has been described that mutations of Rab11 or of the F-BAR / SH3 protein Nervous Wreck, which co-localizes with Rab11 and exerts a role in the negative regulation of BMP by the recycling of its receptors, is associated with a loss of inhibition of this pathway and an increased development of the motor neuron, translated into an increase in the number of synaptic boutons (Khodosh et al., 2006; Rodal et al., 2008). This similarity of morphometric phenotypes would suggest that the loss of inhibition of BMP present in these larvae could be associated with defects in the trafficking of activated receptors in this endosomal recycling compartment.

Taken together, these findings suggest a new mechanism by which the loss of function of an axonal ER structuring protein in the motor neuron could, through its role in the regulation of endosomal trafficking, explain defects in VS accumulation and potentially, the increased signaling of the BMP pathway, even allowing to establish that modifications in membrane trafficking would be part of the etiology of the distal axonopathy present in HSP determined by Atl.

## Author Contribution

Conceptualization: F.B., U.T., J.S.; Methodology: F.B., J.S.; Analysis: F.B.; Investigation: F.B.; Software development resources: B.A, J.J; Resources: F.B., J.S.; Writing & editing: F.B., A.K-S, U.T., J.S.; Supervision: J.S., U.T., C.G.; Funding acquisition: F.B., J.S.

## Acknowledgments

To Dr. Miguel Concha M. for bouton size population analysis, to Dr. Oliver Köbler for introduction to STED microscopy and to Dr. Aaron DiAntonio for dVGLUT antibody.

## METHODS

### Drosophila stocks

Several *Drosophila* strains were used, including: *w*^*1118*^ (Bloomington, USA, BDSC); *wit*^A12^ (BDSC); C380-GAL4 (BDSC); OK6-GAL4 (BDSC), UAS-*dicer2*; UAS-dsRNA-*atl* (Vienna Drosophila Resource Center, Austria, VDRC), UAS-*dicer2*, UAS-*tkv*-CA, UAS-rab4-GFP, UAS-rab11-GFP and Rab7-YFP (BDSC).

### Confocal microscopy

For morphometric analysis, pMAD and synaptic vesicle marker (CSP and VGLUT) fluorescence images were acquired with an Olympus FluoView 1000 confocal microscope, 60X objective **add N.A. 1.35. and oil medium**, 4X zoom and Z step 0.5μm. Images for FM 1-43 analysis were acquired with an Olympus BX61WI Spinning Disc microscope, with a 60X objective **add N.A. 1.42 and oil medium** and 0.5μm Z-step.

### STED microscopy

Images for CSP distribution and colocalization analysis were acquired using a Leica TCS Sp8 STED microscope with a 93X objective **add N.A. 1.3 and glycerol medium**, 5X zoom, reaching 27.5nm resolution in XY and 0.1145μm in Z. STED images were deconvoluted using the Huygens Scripting program (Scientific Volume Imaging, Hilversum, The Netherlands).

### Immunohistochemistry

For larvae body wall dissections and immunostaining protocols were performed as Duncan, Lytle, Zuniga, & Goldstein (2013), respectively. We made some modifications to this protocol, including a shorter fixation time (20 minutes) and a stronger PBT solution (0.3% PBS plus triton X-1000). For STED analysis, the same protocol was used, including differences in the secondary antibodies’ concentrations used. For both confocal and STED microscopy, samples were mounted using the mounting medium Vectashield H-1000 (Vectorlab, USA).

### Antibodies

Primary antibodies: α-DCSP (1:200; Developmental Studies Hybridoma Bank, USA; DHSB), α-pMAD (1:300, MilliporeSigma), α-VGLUT (1:10000, kindly gift from A. DiAntonio, Daniels 2004). Confocal secondary antibodies: Cy5-Hrp y Alexa 594-Hrp, Alexa 488-Hrp, Rhodamine Tritc, FITC and Cy5 α-Mouse or α-Rabbit (1:300, Jackson ImmunoResearch, USA). STED secondary antibodies: Abberior®Ster580 FluoTag-X4-α-GFP, Atto 647N α-Mouse, Atto 594 α-Rabbit (1:200 and 1:300, respectively, Nano-Tag Biotechnologies, Germany), Alexa 594-Hrp; Alexa 488-hrp (1:300, Jackson, USA).

### FM 1-43 assays

Larvae dissection was performed as described in Smith & Taylor (2011). FM 1-43 assay was performed using the protocol described by Gaffield and Betz (2006). Dissected larvae were incubated in 5μM of FM 1-43 (Invitrogen, USA # T35356) for 3 minutes in 90mM KCL in HL3.1 for neuronal stimulation. After stimulation, the larvae were washed 3 times with HL3.1 without calcium and 100 μm of ADVASEP-7 (Biotum, USA #70029) to remove the dye from the surface. Then the larva is incubated in HL3.1 medium and 0.5 mM calcium when images of FM 1-43 loading are acquired. For FM 1-43 unloading, a second chemical stimulation is performed, in absence of FM 1-43 dye addition. Finally, the larva is washed and incubated in HL3.1 with EGTA for the FM 1-43 unloading images.

### Quantification and Statistical analysis

Synaptic bouton number, pMAD and synaptic vesicle markers intensity quantification, as well as FM 1-43 assays were determined in the larvae motor neurons present in the A6 abdominal segment, between muscle 6 and 7. Synaptic bouton quantification was performed using an HRP antibody allowing axonal membrane labeling, while synaptic bouton visualization was attained using α-CSP and α-dVGLUT antibodies. dVGLUT and dCSP synaptic vesicle labels, as well as pMAD label intensity was determined as the average fluorescence intensity of these markers in the synaptic bouton.

We selected the population of synaptic boutons that presented an area size between 2.5-12 μm^2^ to study (type Ib), with exception to the CSP density quartile analysis, in which we quantified boutons of all sizes. This range was determined by measuring the area (μm^2^) of all the synaptic boutons of 10 control larvae in the A6 abdominal segment (muscles 6/7), using α-HRP, to mark the motor neuron membrane and α-CSP for SV. From these data, a size distribution histogram was elaborated and subsequent curve fitting using a Bayesian information criterion (BIC).

Fluorescence intensity quantification in the synaptic bouton was performed as in Andlauer and Sigrist (2012), with slight modifications due to the extended nature of these antibodies labeled inside the bouton (compared to the discrete structures analyzed by these authors). For each independent experiment (at least 2 for each experimental group and driver), all images were acquired using the same optic parameters. ImageJ software (NIH, U.S.A.) was used for image analysis. For each independent experiment, the average of all image backgrounds was subtracted before antibody fluorescence quantification. For fluorescence quantification, using the HRP label we determined the maximal area as a ROI for each synaptic bouton., where synaptic vesicle fluorescence was quantified. These ROIs were used in the VS marker channel, defining the image representing the maximum area of the bouton as the center of the bouton. After obtaining the value of the average intensity of that optical section, the average intensity of the 2 images above and below this section was calculated and added, obtaining a representative sample of the signal contained in each bouton.

ImageJ was also used for FM 1-43 image analysis. For this, from the dye loading images we selected a ROI of each synaptic bouton, using the optical section that represented its maximum area, where we calculated the average fluorescence intensity in each ROI. The average intensity of the Bk of the non-synaptic region located immediately lateral to the bouton was also quantified, which, together with the remote Bk of the image, were averaged and subtracted in each final intensity calculation for each bouton. This calculation makes it possible to rule out that muscle Bk contributes to the final calculation of the average intensity of each bouton. With this method, the load average was calculated, including all quantized synaptic boutons. This same procedure was calculated for the FM 1-43 discharge, where the average fluorescence, calculated from each synaptic bouton present in the discharge image, was divided by the average load of the same larva. This final discharge / load ratio was subtracted from 1, obtaining the discharge percentage. Final values were normalized to control levels within each experiment.

### STED analysis

Program A: Developed in Matlab (Mathworks, USA) by Benedikt Auer, programmer from the LIN institute, with collaboration of Dr. Ulrich Thomas. The program generates a distribution map of the local maxima in the CSP mark, identifying the maximum signal values among neighboring pixels. This program requires the CSP confocal, CSP STED and RAB STED channels of the image file. Briefly, a ROI for each synaptic bouton will be selected from the CSP confocal channel, this will be used as a mask of the respective bouton. Then a Z-stack and the borders of the bouton will be defined. Finally, the CSP STED and RAB STED channel will be selected and analyzed. For each synaptic bouton, two zones are analyzed: A) the peripheral zone, comprising the first 200 nm from the synaptic bouton’s outer edge and where the active zones are located, and B) the central zone, comprising the rest of the inner mark inside the bouton area (Figure 2B) (Bruckner et al., 2012). The program creates binary mask images of the boutons, quantifies the bouton areas and the number of local CSP maxima (hereinafter referred to as CSP particles) and density, and, also, creates a CSP particle distribution map. These were used as input for Program B (see below).

Program B: A macro for the Fiji software (Schindelin et al., 2012) written by Jorge Jara-Wilde (SCIAN-Lab, BNI/U. of Chile) as a post-processing step for the CSP distribution maps and bouton masks generated with Program A. The macro automates the measurement of CSP density variations from the bouton periphery inwards. Starting from the bouton binary image, the total area of the bouton and the CSP density within it are computed. Then, an iterative area shrinking of the bouton region towards its center is implemented by successive morphological erosions. At each iteration, the CSP density is measured within the eroded bouton area. Then, the density of CSP is quantified and accumulated together with the variation of the bouton area, encompassing its entire structure.

Schindelin, J., Arganda-Carreras, I., Frise, E., Kaynig, V., Longair, M., Pietzsch, T., Preibisch, S., Rueden, C., Saalfeld, S., Schmid, B., Tinevez, J. Y., White, D. J., Hartenstein, V., Eliceiri, K., Tomancak, P., & Cardona, A. (2012). Fiji: an open-source platform for biological-image analysis. Nature methods, 9(7), 676–682. doi:10.1038/nmeth.2019

### Colocalization analysis

Colocalization analysis images were acquired at Leibniz-Institut für Neurobiologie (LIN, Magdeburg, Germany), using STED microscopy. For each analysis, the STED channel images for CSP and Rab marker were used, as well as the HRP confocal channel, which allowed delineation of the edge of each synaptic bouton. Using ImageJ, for each synaptic bouton we performed a HRP signal mask, located along the entire Z-axis of each bouton, which was used as ROI to select all the optical slices containing the CSP and Rab information included in that region. In order to determine the colocalization coefficients of both channels, using the JACoP plug-in (Bolte and Cordelières 2006), the CSP and Rab signal were segmented, and the Manders colocalization coefficients M1 and M2 were determined. Here, M1 corresponds to the amount of signal contained in colocalized pixels of the CSP channel divided by its total fluorescence, and M2 corresponds to the same ratio, but for the Rab channel. These coefficient values give an estimate of the percentage of colocalization of one signal over the other (Bolte and Cordelières 2006). This colocalization analysis was performed for CSP with the endosomal markers Rab4, Rab7, and Rab11.

### Statistical analysis

Graphpad Prism 6 software (GraphPad Software, USA) was used for all statistical analysis. Prior to determining the statistical test to be used, a Kolmogorov-Smirnov (KS) normality test was performed. After determining the distribution of the samples analyzed, comparisons between groups were realized using Student’s t-test, Mann-Whitney (MW), parametric ANOVA or Kruskal-Wallis (KW) tests, followed by Tukey or Dunn’s multiple comparisons, respectively. In case of two-way analysis, a two-way ANOVA was performed, followed by Tukey and Sidak post-hoc tests.

## Supplementary figures

**Supplementary Figure S1:**
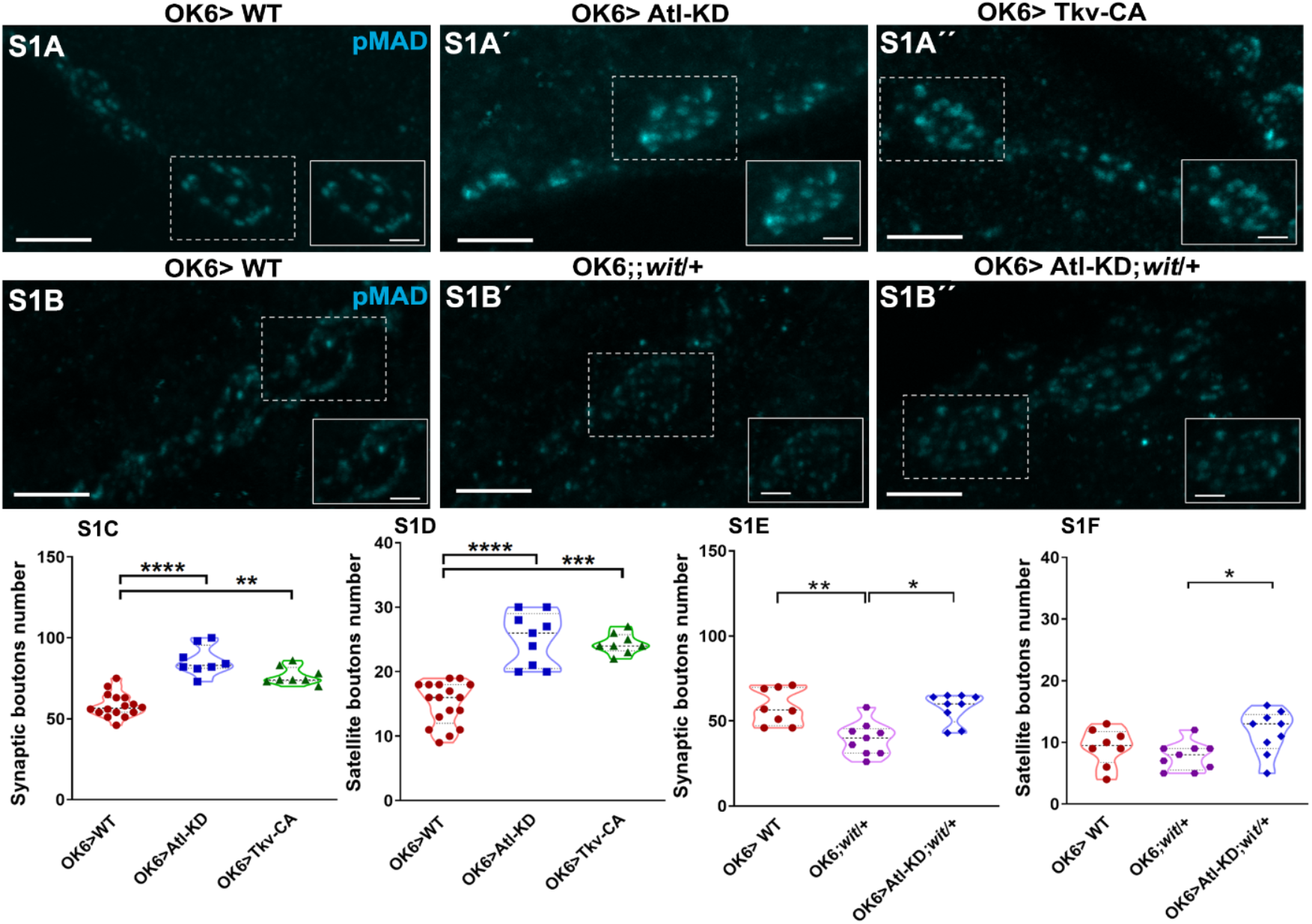
*atl* knock-down in motoneurons (OK6) increases synaptic pMAD and morphometric parameters. A, B) Confocal microscopy images (maximum intensity Z projection) of synaptic pMAD of control (OK6), Atl-KD, constitutive active TkvCA, wit mutant (*wit*) and Atl-KD;*wit* larvae. pMAD antibody stain is labeled in cyan color. Scale bar of large image: 5μm, of cropped image: 2μm. C-D) Synaptic and satellite bouton number of OK6, Atl-KD, TkvCA larvae motor neuron. E-F) Synaptic and satellite bouton number of OK6, *wit* and Atl-KD; *wit* larvae. Each scatter dot represents one measurement. Kruskal-Wallis, p-value *<0,05; **<0.01; ****<0.0001; n=7.

**Supplementary Figure S2:**
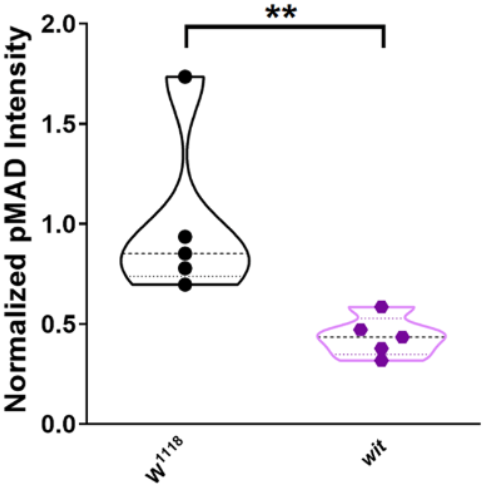
The wit/+ mutant reduces synaptic pMAD in the motor neuron. Synaptic pMAD intensity of w1118 and *wit* mutant larvae. Mann-Whitney, p-value *<0,05; n=5.

**Supplementary Figure S3:**
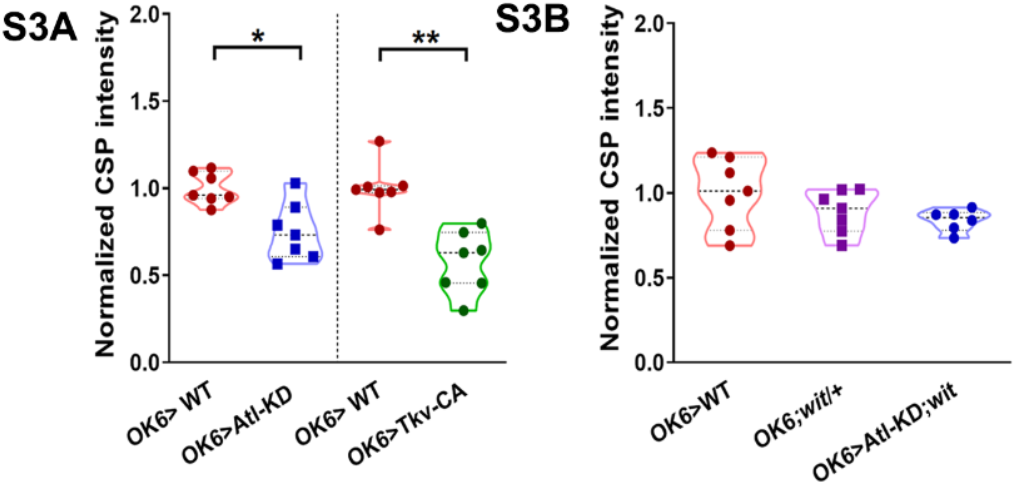
*atl* knockdown reduces **CSP abundance in motor neurons**. S3A) Synaptic CSP intensity of OK6:Atl-KD and TkvCA larvae: CSP intensity normalized to control levels. S3B) Synaptic CSP of OK6; *wit* and Atl-KD; *wit* larvae: CSP intensity normalized to control levels. Each scatter dot represents one measurement. Kruskal-Wallis, p-value *<0,05; **<0.01; n=7.

**Supplementary Figure S4:**
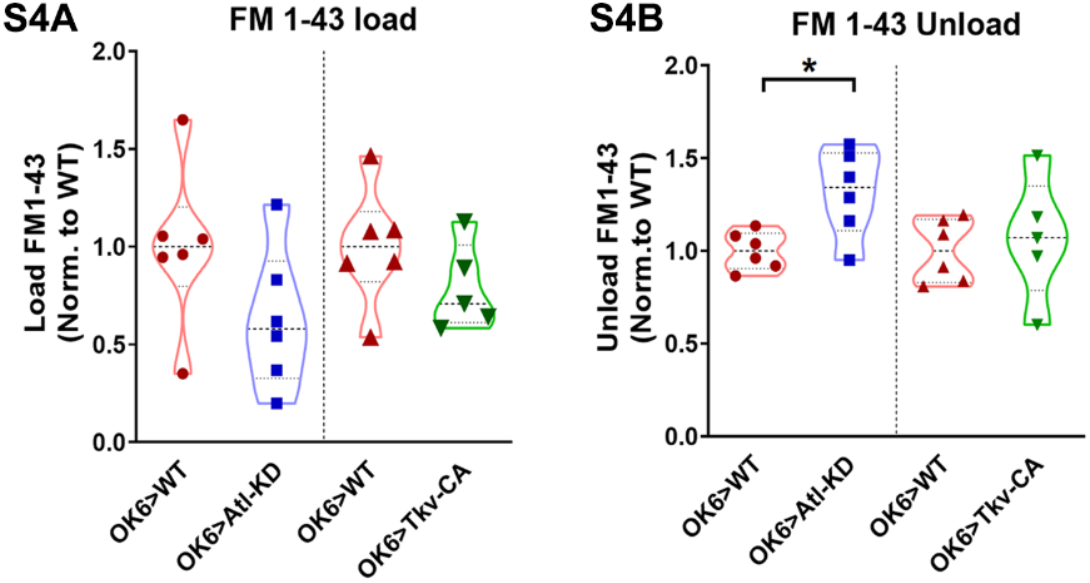
*atl* knock-down in motor neuron increases FM 1-43 discharge. S4A) FM 1-43 loading in OK6, Atl-KD and TkvCA larvae: FM 1-43 load normalized to control levels. S4B) FM 1-43 unloading in OK6, Atl-KD and TkvCA larvae: FM 1-43 unload normalized to control levels. Mann-Whitney, p-value *<0,05; n=7.

